# Detection of cellular microRNAs with programmable DNA nanoswitches

**DOI:** 10.1101/334631

**Authors:** Arun Richard Chandrasekaran, Molly MacIsaac, Paromita Dey, Oksana Levchenko, Lifeng Zhou, Madeline Andres, Bijan K. Dey, Ken Halvorsen

**Affiliations:** The RNA Institute, University at Albany, State University of New York, Albany, NY 12222, USA; Department of Biological Sciences, University at Albany, State University of New York, Albany, NY 12222, USA

**Keywords:** DNA nanoswitch, biosensing, microRNA detection, DNA nanotechnology

## Abstract

MicroRNAs are short non-coding regulatory RNAs that are increasingly used as disease biomarkers. Detection of microRNAs can be arduous and expensive, and often requires amplification, labeling, or radioactive probes. Here we report a single-step, non-enzymatic detection assay using conformationally responsive DNA nanoswitches. Termed miRacles (<u>mi</u>cro<u>R</u>NA <u>a</u>ctivated <u>c</u>onditional <u>l</u>ooping of <u>e</u>ngineered <u>s</u>witches), our assay has sub-attomole sensitivity and single-nucleotide specificity using an agarose gel electrophoresis readout. We detect cellular microRNAs from nanogram-scale RNA extracts of differentiating muscle cells, and demonstrate multiplexed detection of several microRNAs from one biological sample. We demonstrate one-hour detection without expensive equipment or reagents, making this assay a compelling alternative to qPCR and Northern blotting.

**Significance statement:** Detection of microRNAs play a key role in biological research and medical diagnostics, and current detection methods are expensive and require sophisticated processes. We present <u>mi</u>cro<u>R</u>NA <u>a</u>ctivated <u>c</u>onditional <u>l</u>ooping of <u>e</u>ngineered <u>s</u>witches (miRacles), a mix-and-read strategy that is based on conformational changes of DNA nanoswitches upon binding a target microRNA. MiRacles has a sensitivity of ∼4 copies/cell and specificity of a single nucleotide, and can be performed in one hour at a fraction of the cost of traditional microRNA detection techniques. Our method can also be multiplexed to detect multiple microRNAs from one biological sample. The minimalistic miRacles assay has immediate application in biomedical research and longer term potential as a clinical tool.

## Main Text

MicroRNAs are short, noncoding RNAs (18-25 nt) that repress gene expression at the post-transcriptional level. They impact many biological processes including cellular proliferation, differentiation and apoptosis, leading to important consequences in normal development, physiology, and disease [1-2]. Expression levels of individual microRNAs in tissues, cells, and bodily fluids can serve as stable biomarkers for cellular events or disease diagnosis [3-4], highlighting the importance of a simple and sensitive method for microRNA detection and quantification.

MicroRNA detection is challenging due to low abundance, small size, and sequence similarities. MicroRNAs comprise about 0.01% of total RNA [5], and individual microRNA levels range from a few copies to tens of thousands of copies per cell [6]. Furthermore, microRNAs within a family can differ by a single nucleotide [7], and yet each specific microRNA can be differentially regulated during cellular processes or in disease conditions. Thus, microRNA detection strategies must be highly specific, able to correctly identify a few target molecules among an abundance of similar RNA molecules.

Traditional methods for detection of microRNAs include Northern blotting, quantitative real-time PCR (qRT-PCR), next generation sequencing and microarray-based hybridization [5,8,9]. Of these, only Northern blotting detects native microRNA directly, while the others rely on additional labeling or amplification steps. These methods tend to have significant tradeoffs between cost and performance, typically require skilled personnel, involve complex and time consuming procedures, and require specialized equipment.

Here we introduce the miRacles assay: <u>mi</u>cro<u>R</u>NA <u>a</u>ctivated <u>c</u>onditional <u>l</u>ooping of <u>e</u>ngineered <u>s</u>witches. The miRacles assay uses a “smart reagent” comprised of rationally designed DNA nanoswitches to enable simple and low-cost detection of native microRNAs without specialized equipment. As we show here, our assay can be used to detect and quantify microRNAs from nanogram-scale RNA extracts, and can be performed in as little as an hour with common lab supplies.

## Results

The DNA nanoswitch [10-11] was designed as a linear duplex that forms a loop in the presence of a target microRNA (**Figure 1A and Supplementary Figures S1-S2**). The nanoswitch was constructed using DNA origami approaches [12], formed by hybridizing short oligonucleotides (typically 60 nt) that are complementary to a single-stranded DNA scaffold (7249 nt). Two distant “detector” strands (separated by ∼2500 nt) were designed to contain overhangs complementary to different segments (typically halves) of the target microRNA. Recognition and binding of the microRNA reconfigures the switch from the linear “off” state to the looped “on” state. The two states can be quantified using standard agarose gel electrophoresis and gel stains (**Figure 1B**), where the detection signal arises from the integrated intensity of the looped nanoswitch scaffold DNA. Each nanoswitch recruits thousands of intercalating dye molecules, but has its fate (looped or unlooped) decided by a single microRNA, providing an inherent signal amplification.

**Figure 1.**
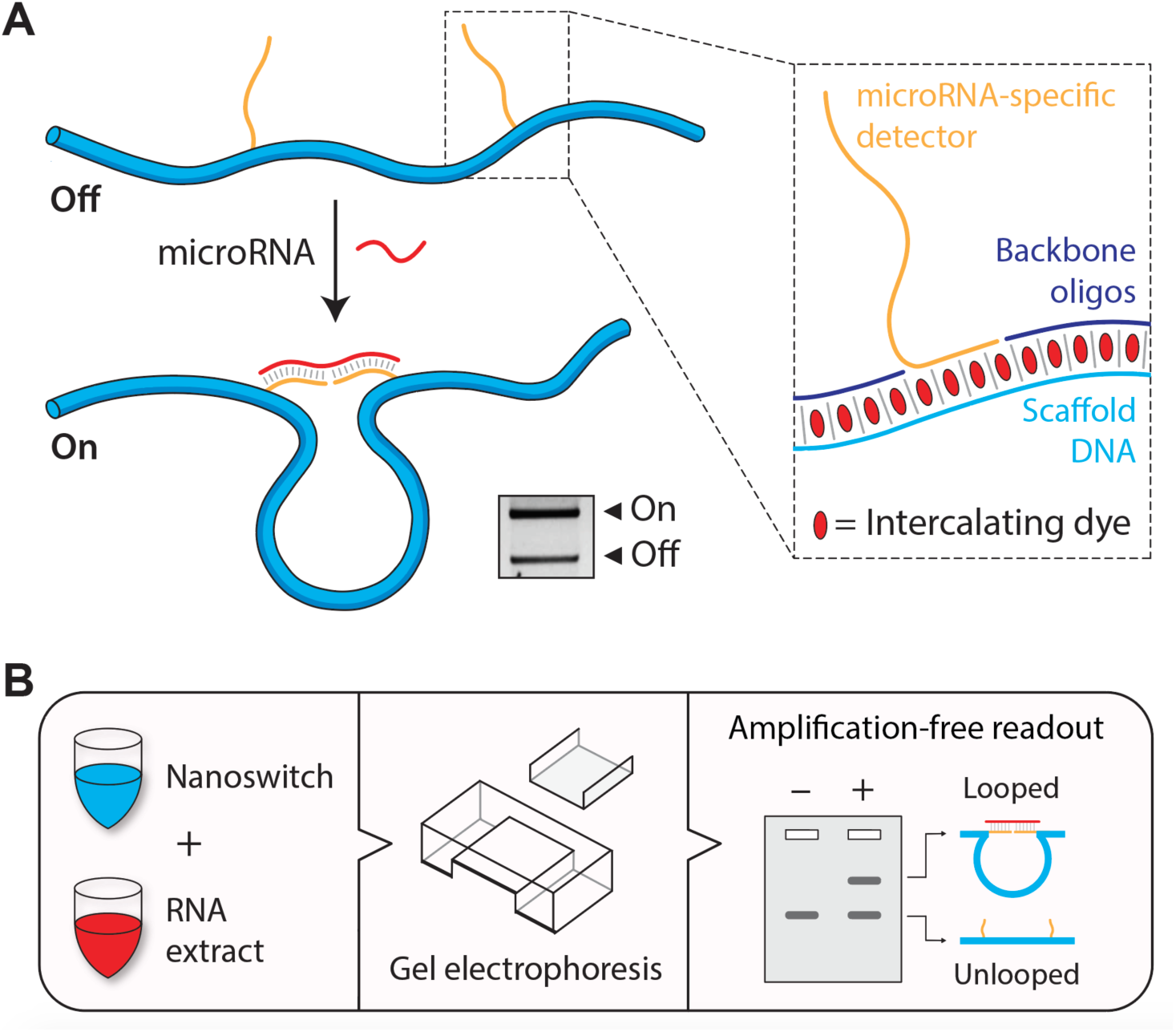
Concept and workflow of the miRacles assay. (A) DNA nanoswitches undergo a conformational change from a linear ‘off’ state to a looped ‘on’ state when bound to a target microRNA. The two conformations are resolvable in a standard agarose gel. Inset: The nanoswitch is composed of a single stranded M13 scaffold, backbone oligonucleotides and single stranded extensions (detectors) complementary to the target microRNA. (B) Workflow of the **miRacles** assay: customized DNA nanoswitches are mixed with target microRNA sample, incubated and run on a agarose gel for detection.

For concept validation, we chose let-7b as a target microRNA, because let-7b belongs to a highly conserved family of over a dozen related microRNAs varying by one or more nucleotides. These microRNAs have critical biological functions and are dysregulated in multiple human diseases [13]. We customized DNA nanoswitches with detector strands that target the full sequence of let-7b, and confirmed microRNA detection by incubating them with synthetic let-7b microRNA and running an agarose gel (**Figure 1A, gel inset**).

Next, we investigated the ability of our let-7b nanoswitches to distinguish closely related sequences by testing either synthetic let-7c target (1-nt mismatch) or synthetic let-7a target (2-nt mismatch). We found that a one-nucleotide mismatch in let-7c caused an 85% reduction in signal intensity compared to let-7b. Interestingly, only a two-nucleotide mismatch in let-7a completely abolished the signal (**Supplementary Figure S3**). However, we found that reducing the detector length on the mismatch side completely eliminated the crosstalk signal in the one-nucleotide mismatch (**Figure 2A**). These dramatic results illustrate the high specificity of our assay, which has been a key challenge for microRNA detection [8-9].

**Figure 2.**
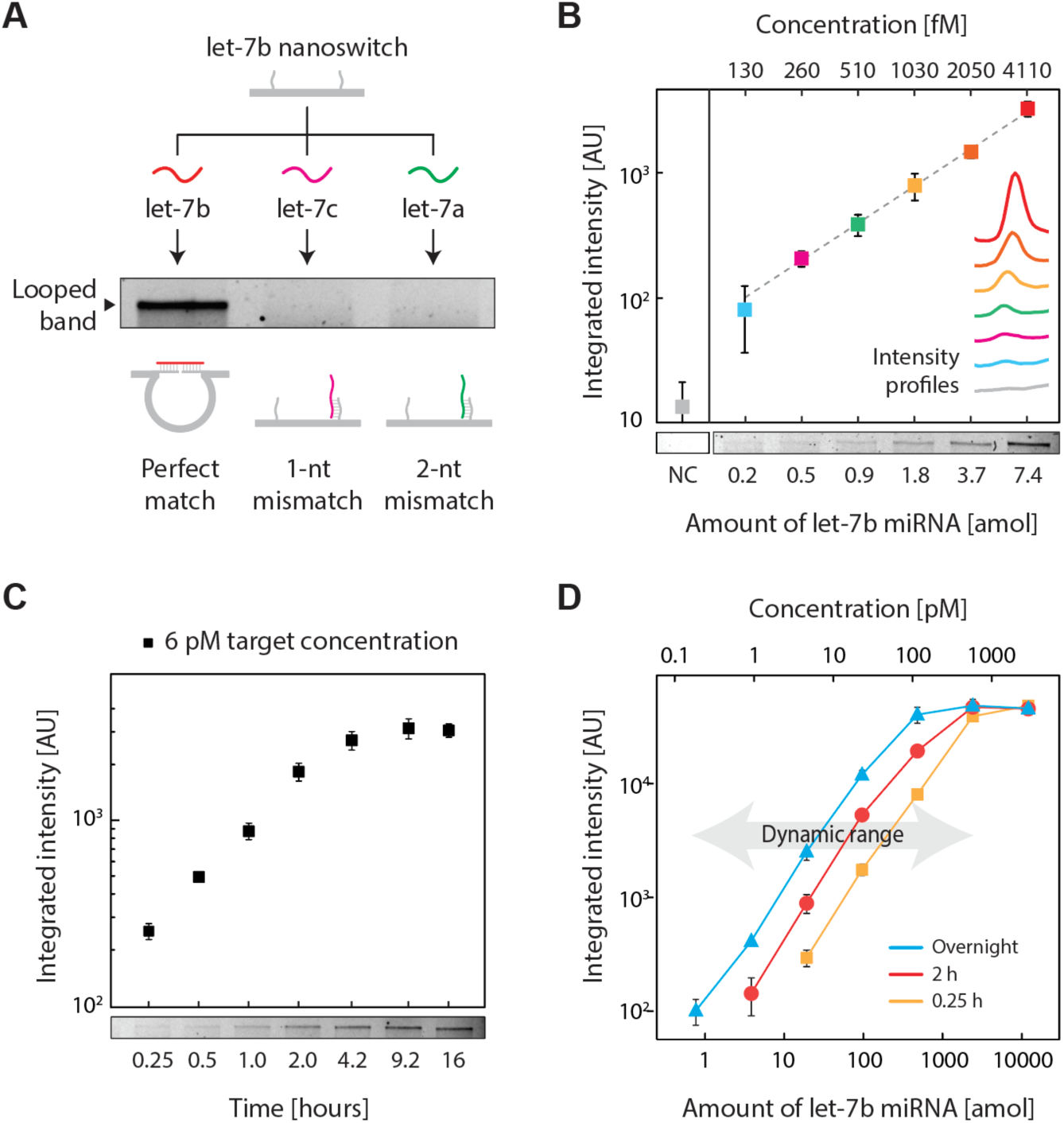
Validation of the miRacles assay. (A) Specificity of the DNA nanoswitches with detectors designed for let-7b. As low as 1-nt mismatch between the detectors and the target microRNA eliminates the signal. (B) Limit of detection of the assay, (C) Time course of the assay for a low concentration target, (D) Dynamic range of the assay at different reaction times. Data points and error bars represent the mean and standard deviation, respectively, from triplicate measurements.

The often-low abundance of microRNA requires high-sensitivity detection. We show a limit of detection of ∼ 0.2 attomoles (**Figure 2B and Supplementary Figure S4**), corresponding to ∼100,000 molecules. We measured the time course for detection of a low concentration target (6 pM), which shows increasing signal until ∼4 hours with little change beyond that time (**Figure 2C and Supplementary Figure S5**). To explore the full dynamic range, we used various incubation times to demonstrate microRNA detection spanning over 4 orders of magnitude (**Figure 2D and Supplementary Figure S6**), which closely mirrors the natural dynamic range of different microRNAs [6].

Moving beyond synthetic targets, we sought to establish the miRacles assay in cellular RNA extracts. MicroRNAs are commonly studied in cell cultures, tissue samples, and body fluids, all of which can be readily processed to collect total RNA (typically microgram scale) and a small RNA subfraction (typically ∼10% by mass). For this work, we used myoblast cells as a surrogate model system for muscle differentiation and miR-206 as our primary target. Among the first identified and most important microRNAs in differentiating skeletal muscle cells, miR-206 undergoes significant upregulation, especially in late differentiation [14-15]. We harvested undifferentiated and differentiated muscle cells and extracted total and small RNAs from these cells (**Figure 3A**). We confirmed differentiation in these cells by Western blots (**Figure 3B**) and immunocytochemistry (**Figure 3C**) of early and late myogenic differentiation markers myogenin (Myog) and Myosin Heavy Chain (MHC), respectively.

**Figure 3.**
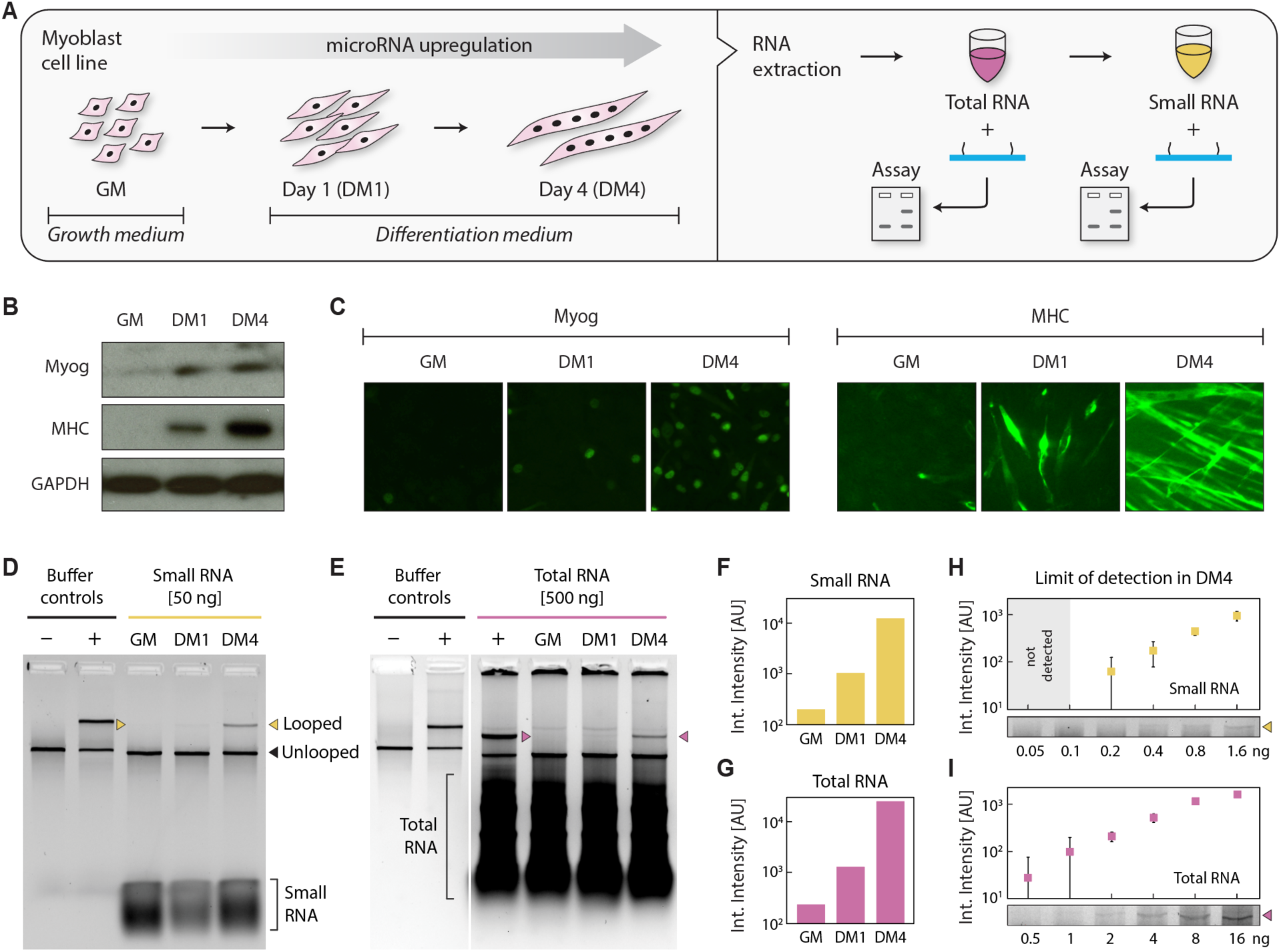
MicroRNA detection from differentiating myoblast cells. (A) Schematic showing myoblast cells, harvested while growing in GM and on differentiation days 1 and 4, processed to yield total and small RNA fractions. An early myogenic differentiation marker, myogenin (Myog) and a late myogenic differentiation marker, Myosin Heavy Chain (MHC) were measured by (B) Western blotting and (C) by immunocytochemistry to confirm differentiation. Both Myog and MHC were upregulated in DM1 and DM4. GAPDH served as a control in (B). (D) We detected miR-206 in the differentiated samples with 50 ng of small RNA and (E) with 500 ng of total RNA. Quantification of gel intensities shows a sharp progressive upregulation during differentiation, similar in both (F) small RNA and (G) total RNA samples. From DM4 samples, we note detection from as little as (H) 200 pg small RNA and (I) 500 pg total RNA. Error bars represent standard deviation from triplicate measurements.

Individual microRNA species are commonly at the parts per million level by mass of total RNA, illustrating the “needle in a haystack” type challenge faced in microRNA detection. To validate our assay for biological detection, we first started with the less complex small RNA subfraction. Using miR-206 targeting nanoswitches, we observed a progressive increase in miR-206 expression in small RNA between undifferentiated samples and differentiation days 1 and 4 (**Figure 3D, 3F**). This sharp upregulation is similar to previous observations by qRT-PCR [15]. We further showed that by altering a single nucleotide on one of the detector strands, the detection signal was eliminated (**Supplementary Figure S7**). In the less processed but more complex total RNA samples, minor protocol modifications enabled miR-206 detection clearly in 500 ng, with a similar trend to the small RNA fraction (**Figure 3E, 3G & Supplementary Figure S8**). Focusing on the highly expressed miR-206 in the late differentiation samples (DM4), we detected miR-206 in as little as 200 pg of small RNA and as little as 500 pg of total RNA (**Figure 3H, 3I & Supplementary Figure S9**). Assuming ∼20 pg of total RNA/cell, this corresponds to only 25 cells. Given our established limit of detection of 0.2 attomole (∼100,000 molecules) these results suggest that miR-206 is present in ∼4000 copies per cell in the DM4 samples.

Often, multiple microRNAs alter their expression level during different cellular or disease stages. We used the the programmability of the nanoswitches to develop a multiplexing system capable of detecting multiple microRNAs from the same sample. We chose 4 microRNAs known to be present in these muscle cells (miR-206, miR-125b, miR-24, and miR-133-3p), and 1 negative control microRNA (miR-39) specific to *Caenorhabditis elegans*. We designed 5 individual nanoswitches with different loop sizes and combined them to form a multiplex system that produces a distinct band for each target sequence (**Figure 4A**). In 50 ng of small RNA, we detected the four microRNAs at various expression levels spanning nearly two orders of magnitude, and showed no detection of the negative control (**Figure 4B**). This multiplexing strategy enables direct comparison of microRNA levels in one sample without labeling or amplification, and also moves a step in the direction of expanding the throughput of the miRacles assay.

**Figure 4.**
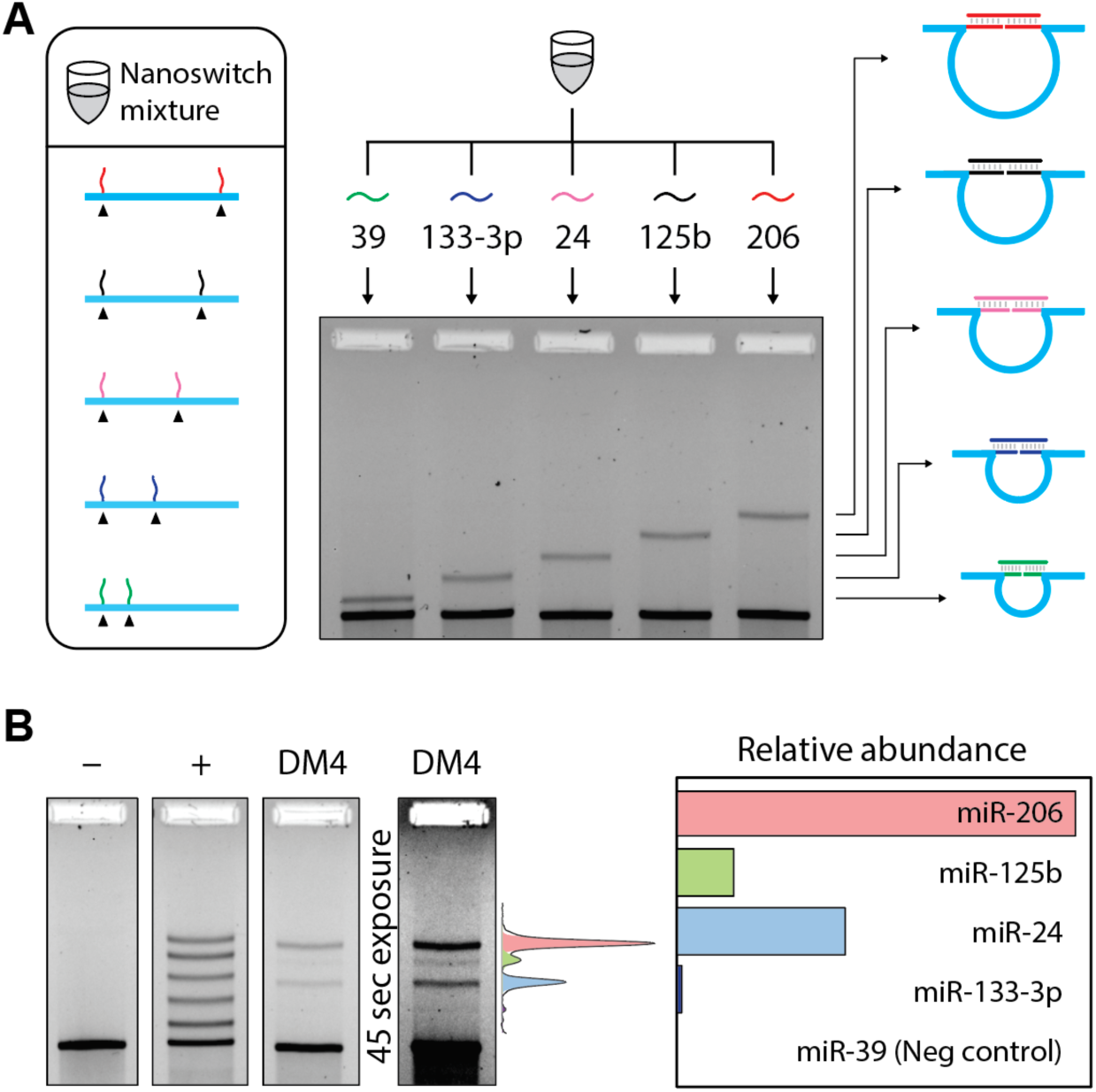
Five-channel multiplexing. (A) Multiplexing enables the detection of different microRNAs with different loop sizes. (B) A multiplexed nanoswitch mixture shows 5 bands with similar intensity in a positive control consisting of all 5 target microRNAs. In 50 ng of DM4 small RNA, 4 different microRNAs are detected at various expression levels, with miR-39 (a *C. Elegans* specific microRNA) not being detected.

## Discussion

Performance of the miRacles assay is competitive with commonly used techniques (**Supplementary Table 1**). Our selectivity of 1 nt has been difficult to achieve using other methods [8-9], and our sensitivity outperforms non-amplified methods including Northern blotting and microarray. While we cannot match the theoretical single-molecule sensitivity of qRT-PCR, in practice such sensitivity is rarely achieved or required. Our 0.2 amol limit of detection (∼100,000 molecules) suggests that a 500 ng sample of total RNA (∼25,000 cells) would enable detection of ∼4 copies per cell. Given this low detection level, increased sensitivity is not especially meaningful for most cellular extracts. Since our assay directly measures microRNA without needing amplification, protocols are simpler and avoid extra sample processing that can introduce errors. Uniquely, our assay enables multiplexed quantitation of multiple native microRNAs from a single sample.

In stark contrast to other techniques, our minimalistic approach to microRNA detection is most notable for what is absent: there is no labeling, no amplification, no surface binding, no wash steps, no expensive equipment or reagents, no complex protocols. Our DNA nanoswitches can be custom-made in advance at a cost of less than one penny per reaction (**Supplementary Table S2**) and stored (wet or dry) for long-term use (**Supplementary Figure S10**). Once made, simply mixing the nanoswitches with the sample liquid and running a gel produces high quality results with common lab supplies. We illustrate this with a time-lapse video of an undergraduate researcher performing start-to-finish microRNA detection from a biological sample in 1 hour (**Supplementary Video S1**).

Our assay is immediately useful for measuring microRNAs from biological samples, but also has potential for clinical applications. Detection of microRNA biomarkers [3-4] and biomarker panels [16] from bodily fluids is increasingly relevant for diagnosing and monitoring diseases, and DNA nanoswitches have already shown compatibility with serum and urine [11,17]. Scalable production with liquid handlers and liquid chromatography based purification [18] could generate a complete library of microRNA nanoswitches, stored without refrigeration and used with minimal laboratory infrastructure. With these features in mind, we align with the broader concept of frugal science [19], a movement that has already produced clever low-cost solutions to blood centrifugation [20] and water purification [21], among others. As with those examples, we disrupt the typical cost/performance relationship, bringing simple microRNA detection to everyone’s fingertips with mix-and-read smart reagents.

## Acknowledgments

We thank M. Belfort, P. Rangan, G. Fuchs, S. Ranganathan, and W.P. Wong for critical comments and suggestions on the manuscript. Research reported in this publication was supported by the NIH through NIGMS and NCI under awards R35GM124720 and R21CA212827, respectively to K.H. The content is solely the responsibility of the authors and does not necessarily represent the official views of the National Institutes of Health.

## Author Contributions

A.R.C., B.K.D., and K.H. designed experiments; A.R.C., M.M., O.L., L.Z., M.A., and K.H. performed nanoswitch experiments; P.D. and B.K.D. carried out cell culture, myoblast differentiation, total and small RNA extraction from these samples and performed western blot and immunocytochemistry experiments. A.R.C., B.K.D., and K.H. wrote the manuscript, all authors provided comments and took part in editing.

